# Protlego: A Python package for the analysis and design of chimeric proteins

**DOI:** 10.1101/2020.10.04.325555

**Authors:** Noelia Ferruz, Jakob Noske, Birte Höcker

## Abstract

**Motivation:** Gene duplication and recombination of protein fragments have led to the highly diverse protein space that we observe today. By mimicking this natural process, the design of protein chimeras via fragment recombination has proven experimentally successful and has opened a new era for the design of customizable proteins. The *in-silico* building of structural models for these chimeric proteins, however, remains a manual task that requires a considerable degree of expertise and is not amenable for high-throughput studies. Energetic and structural analysis of the designed proteins often require the use of several tools, each with their unique technical difficulties and available in different programming languages or web servers.

**Results:** We have implemented a Python package that enables automated, high-throughput design of chimeras and their structural analysis. First, it is possible to fetch evolutionarily conserved fragments from a built-in database (also available at fuzzle.uni-bayreuth.de). These relationships can then be represented via networks or further selected for chimera construction via recombination. Designed chimeras or natural proteins are then scored and minimised with the Charmm and Amber forcefields and their diverse structural features can be analysed at ease. Here, we showcase Protlego’s pipeline by exploring the relationships between the P-loop and Rossmann superfolds and building and characterising their offspring chimeras. We believe that Protlego provides a powerful new tool for the protein design community.

**Availability and implementation:** Protlego is freely available at (https://hoecker-lab.github.io/protlego/) with tutorials and documentation.

## 1 Introduction

Proteins evolved to form diverse structures and perform a multitude of functions. If we could unravel basic rules to design new proteins and implement tailored functions, this could address many challenges of today’s society. The design of customized proteins, however, has not been an easy task. There are by now an impressive set of successful examples for *de novo* designed protein structures (Kuhlman *et al*., 2003; Huang *et al*., 2014, 2016; Thomson *et al*., 2014). However, the majority of engineered enzymes are still obtained via directed evolution starting from a natural protein (Lechner *et al*., 2018). Also the recombination or duplication of natural protein segments has led to new proteins (Höcker, 2014). Interestingly, nature seems to have created the vast protein space we observe today by these latter mechanisms, i.e. via the duplication and recombination of protein domains. Domain recombination has led to the development of large multidomain proteins, whose synergic effects enable differentiation and speciation of functionalities (Dohmen *et al*., 2020). Domains are allegedly the basic evolutionary unit (Apic and Russell, 2010; Ponting and Russell, 2002), and significant efforts have been made to hierarchically classify them, such as in the SCOP (Fox *et al*., 2014), CATH (Dawson *et al*., 2017), and ECOD (Cheng *et al*., 2014) databases. However, the origin of domains themselves is known today to derive from the duplication, recombination and differentiation of sub-domain sized fragments (Lupas *et al*., 2001; Söding and Lupas, 2003).

We could show that two major protein folds, the TIM-barrel and the flavodoxin-like fold, are evolutionarily related and share a fragment of common origin (Farías-Rico *et al*., 2014). Further, Alva *et al* identified a set of 40 peptides of up to 38 amino acids in length whose sequence similarity is evidence of common ancestry despite appearing in different folds (Alva *et al*., 2015). Similarly, we recently performed an all-against-all comparison of protein domains representing all existing folds and identified more than 1000 conserved protein fragments of various lengths across protein space. These fragments are present in different protein environments and represent building blocks that nature has reused through the course of evolution and that can now be browsed in the publicly available Fuzzle database (www.fuzzle.uni-bayreuth.de)(Ferruz *et al*., 2020).

Engineering efforts have been successful in designing new protein domains by duplicating or recombining protein fragments. The design of symmetric protein structures through duplication of the same fragment has been achieved in the design of TIM-barrels (Höcker *et al*., 2004; Fortenberry *et al*., 2011), β-trefoils (Lee and Blaber, 2011), β-propellers (Yadid and Tawfik, 2011; Voet *et al*., 2014), TPR-proteins (Zhu *et al*., 2016), Armadillo-repeat proteins (Reichen *et al*., 2016; Parmeggiani *et al*., 2015), and outer membrane β-barrels (Arnold *et al*., 2007). The design via recombination of fragments within as well as across folds has also been effective (Höcker, 2014). In a set of two pioneering studies, Riechmann and Winter obtained folded proteins through recombination of random polypeptide segments (Riechmann and Winter, 2000, 2006). It was further possible to design a chimeric TIM-barrel through the combination and optimization of HisF, a TIM-barrel protein, and CheY, a flavodoxin-like domain (Bharat *et al*., 2008; Eisenbeis *et al*., 2012). In a follow-up study the combination of HisF with another flavodoxin, NarL, led to a protein of even higher stability (Shanmugaratnam *et al*., 2012).

While the design via recombination is fairly new, its success provides an alternative to classical design approaches. Individual fragments can contribute their unique functional properties to the chimeric protein, which provides an interesting, alternate, and generalizable route for protein design. However, the *in silico* automated design of chimeric proteins as a prior step for experimental studies requires broad expertise and the use of several tools (Farias-Rico and Höcker, 2013).

A few algorithms have been made available for the recombination of homologous sequences. In 2002, Frances Arnold and her group published SCHEMA, an algorithm to detect segments of homologous proteins that can be recombined without disturbing the integrity of the structures. The resulting sequences produce folded hybrid proteins with a greater likelihood than by random shuffling (Voigt *et al*., 2002; Meyer *et al*., 2003). The authors later described an algorithm for the creation of chimera libraries that are enriched in folded proteins without compromising the diversity (RASPP) (Endelman *et al*., 2004). The software is available at https://github.com/mattasmith/SCHEMA-RASPP and confers a useful tool for the creation of recombination libraries from homologous parents. In addition the software MODELLER provides a useful software for the modelling of protein structures based on a sequence alignment (Sánchez and Sali, 2000), and thus has been valuable for specific scenarios with a few chimeras (Farias-Rico and Höcker, 2013). Other well-known similarly working homology modelling predictors are SwissModel (Peitsch, 1995), FAMS (Ogata and Umeyama, 2000) and EsyPred3D (Lambert *et al*., 2002). More recently, machine learning methods have led to higher-accuracy predictors, such as DMPfold (Greener *et al*., 2019) or AlphaFold (Senior *et al*., 2020). On a recent study, the Kuhlman lab defined a new computational approach, termed SEWING, which enabled the creation of chimeric all-alpha structures by joining short alpha helical fragments (Jacobs *et al*., 2016) and optimizing the chimeras with the program ROSETTA (Leaver-Fay *et al*., 2011). The method was experimentally tested, and the resulting X-ray structures matched the predictions. To our knowledge, the method has not been applied to the construction of non all-alpha chimeras though.

Here, we implemented an easy-to-use python package named Protlego that automates the process of *in silico* chimera design and structural analysis. Concretely, the software is divided into five main applications (**Fig. 1**). Protlego contains a light built-in Fuzzle database, comprising more than 10 million entries of evolutionary related protein fragments that appear in different protein contexts. Upon selection of one or more fragments from the database, evolutionary relationships can be represented via networks or offspring chimeras can be built from two protein parents in an automated fashion. The resulting chimeras can be directly ranked by scoring or minimizing with the molecular mechanics forcefields CHARMM or Amber. We have implemented a set of tools that enable the structural analysis and energetic evaluation of the chimeras or any PDB of the user’s choice. As an example, one can visualize hydrogen bond and salt bridge networks, as well as hydrophobic clusters in the protein structure.

**Figure 1:**
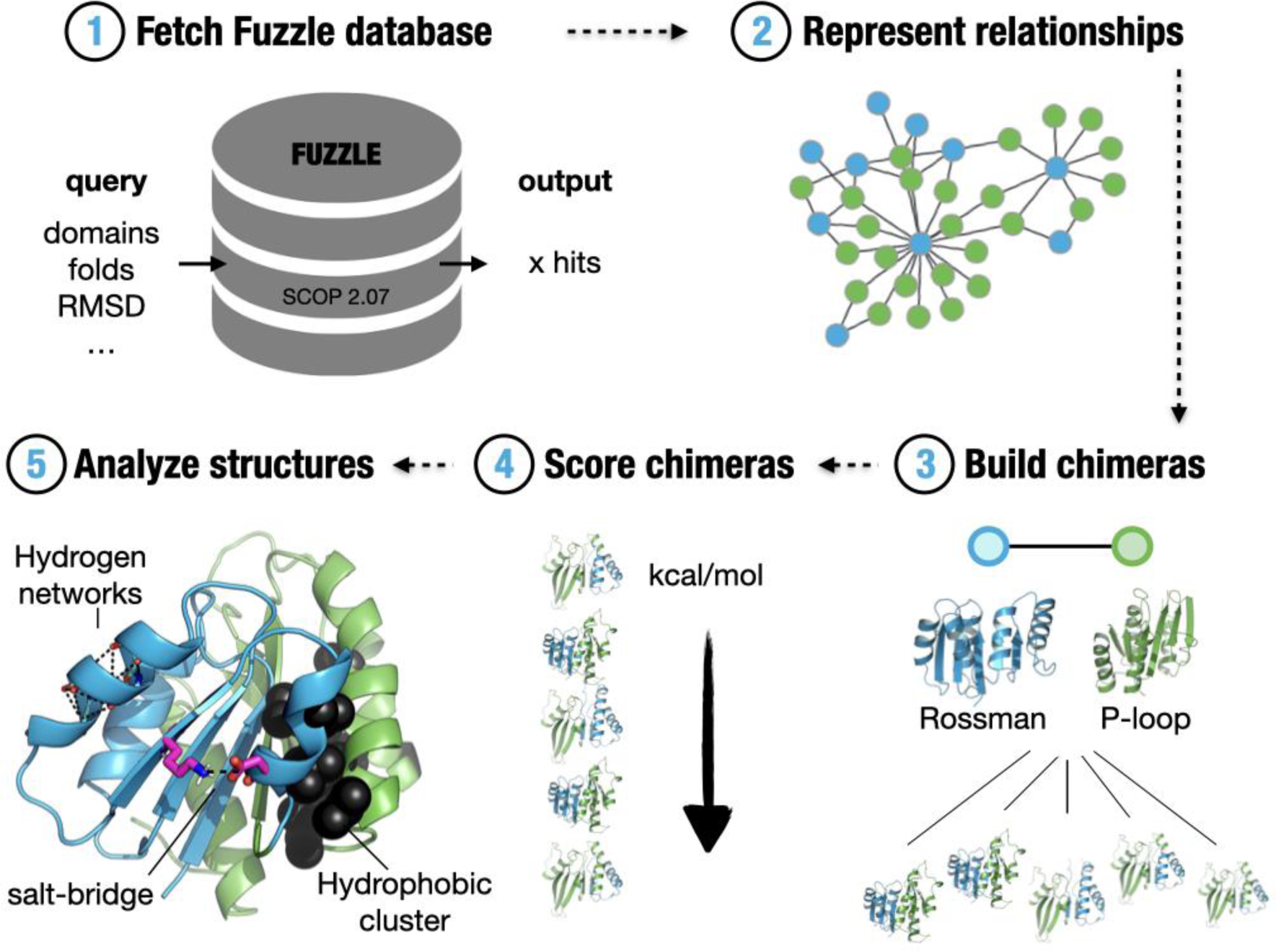
Overview of Protlego features. Protlego is mainly divided into five applications: It is possible to fetch from a built-in Fuzzle database (1) and represent the hits via similarity networks. The hits can also be used to build chimeras (3), which can later be scored (4) and analysed (5)

Here, we showcase Protlego’s features by exploring the relationships between two α/β superfolds, namely, the P-loop containing nucleoside triphosphate hydrolases (NTPases) and the Rossmann fold. Tutorials to reproduce these exact results and guidelines to customize different analysis are freely available at https://hoecker-lab.github.io/protlego/ and **Listing S1**. We believe that Protlego will be a useful tool for protein engineers and evolutionary biologists alike.

## 2. Methods

### 2.1 Fetching hits from the Fuzzle database

Protlego contains a lightweight Fuzzle database that gets installed during setup via sqlite3. The Fuzzle database was created by doing an all-against-all profile hidden Markov model (HMM) comparison of all domains in SCOPe95 2.07 (Fox *et al*., 2014) using HHsearch (Söding, 2005). Fuzzle contains more than 10 million hits among over 28 000 unique domains (**Fig. 1,1**). Each hit in Fuzzle contains information about the two domains that contain a common fragment (denoted *query* and *subject*), start and end of the fragment they share, HHsearch probability, and RMSD, among others (Ferruz *et al*., 2020). Protlego enables fetching hits from the Fuzzle database searching via PDB identifier, domain identifiers, specific SCOPe groups (families, superfamilies and folds). It is also possible to fetch entire subspaces that fulfil certain criteria (such as RMSD or length below a certain threshold, to name a few examples). An overview of the methods available in this application is presented in **Table 1**.

**Table 1:** Fetching functions to retrieve from the built-in Fuzzle database.

| Fetching function | Hits that the function fetches from Fuzzle |
| --- | --- |
| fetch_id | Hit with that Fuzzle ID |
| fetch_by_domain | Hits that contain a specific domain as query or subject |
| fetch_by_domains | Hits between the two domains |
| fetch_by_PDB | Hits that contains domains that belong to the PDB |
| fetch_by_PDBs | Hits between two domains that belong to PDB1 and PDB2 |
| fetch_group | Hits belonging to one or hits between two specific SCOPe groups (folds, superfamilies, families). |
| Fetch_subspace | Hits that satisfy the cut-offs for RMSD, length, and TM-score, among others. |

All include RMSD, minimum and maximum lengths, or TM-score as optional parameters to refine the search.

### 2.2 Network visualization

Protlego facilitates the visualization of evolutionary relationships via similarity networks (**Fig. 1,2**). The Graph class takes a hit or set of hits from the built-in database and enables its representation harnessing the power of the *graph-tool* package (Peixoto T, 2014). Network nodes represent protein domains and links join domains that have a fragment in common. It is possible to directly visualize nodes (the fragment in the context of its domain) and links (the alignments between two domains that share a fragment) using VMD.

### 2.3 Chimera modelling

Protlego builds all possible chimeras between two protein parents (**Fig. 1,3** and **Fig. 2**). The chimeras are built by combining N- and C-terminus from query and subject, and thus present a single recombination point. All possible chimeras from the two combinations are built, where *combination1* refers to those chimeras where the N-terminus comes from the query, and *combination2* to those where it comes from the subject. The Builder class takes as an argument a hit and uses the HHsearch alignment as a template to create the models. The aminoacids in the local alignment get mapped to their corresponding alpha carbon atoms in the PDB structures (**Fig. 2a**). The alignment of the PDB structures is performed with TMalign (Zhang and Skolnick, 2005) taking only into account the fragment’s Cα atoms. There are two ways to perform this alignment, either by minimizing the RMSD taking all the alpha carbons into account in a global fashion, or by performing stepwise partial alignments. In the case of a long fragment, the partial mode iteratively finds the best alignments for shorter regions such as, for example, βα motifs. These regions are defined by the sequence alignment: sequence alignment gaps constitute the boundaries that define each shorter region.

**Figure 2:**
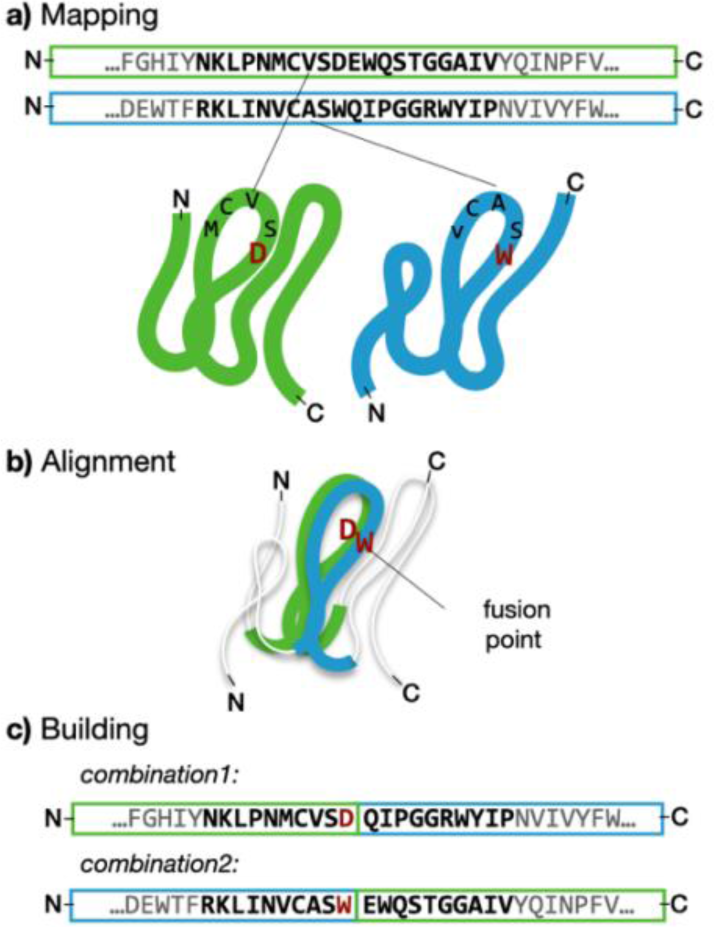
Summary of the chimera building process in Protlego. First, the aminoacids in the HHsearch alignment are mapped onto the corresponding PDB structures **(a)**. These structures are then superimposed by minimizing the RMSD of each pair of alpha carbons with TMalign **(b)**. At this stage, distances for each fusion point are computed and each point below a cut-off is defined as a fusion point. Each fusion point can produce 2 chimeras, one with the query N-terminus (*combination1*) and the other with the subject N-terminus (*combination2*) **(c)**. The process is repeated for each fusion point. Final chimeras are accepted if they do not present backbone clashes.

The rationale behind the partial alignment is to obtain better quality local alignments that will lead to more fusion points. Once the alignment is performed, Protlego computes the distance of each pair of aligned alpha carbons (**Fig. 2b**). Low distances, sometimes found in secondary structure elements, confer ideal fusion points for recombination, as the merging of the two parents ensures that the resulting structure will be minimally perturbed. We use the default value of 1 Å as a maximum distance, albeit users can define their own thresholds. Once the optimal fusion points are found, we recombine query and subjects at these positions (**Fig. 2c**). Only chimeras that do not present backbone clashes are kept. The handling of PDBs is performed with the *moleculekit* package (Doerr *et al*., 2016).

### 2.4 Potential energy evaluation

Protlego allows the estimation of the potential energy of Chimera objects with different versions of the CHARMM (Vanommeslaeghe *et al*., 2009) and Amber (Ponder and Case, 2003) forcefields (**Fig. 1d**). The structures can be scored or minimized accounting the protein backbone when desired. GPU acceleration is supported. The potential energy estimation uses functionality of the *openMM* package (Eastman *et al*., 2017).

### 2.5 Structural analysis

Protlego enables the computation of several structural features for the designed chimeras or natural proteins (**Fig. 1,5**). The user can automatically fetch a PDB or SCOPe domain within the Chimera class. Among others, the computation of solvent accessible surface area (SASA), distance matrices, contact orders, contact maps, hydrophobic clusters and *HHbond* plots (Bikadi *et al*., 2007) is implemented. The automatic visualization of structural features in the protein context is possible. The computation of hydrogen networks utilizes some of the functionality of the *MDtraj* package (McGibbon *et al*., 2015). Protlego also includes an implementation of the CSU algorithm (Sobolev *et al*., 1999) which enables the computation of hydrophobic clusters.

#### 2.5.1 Computation of hydrophobic clusters

It has been proposed that sidechains of isoleucine (ILE), leucine (LEU) and valine (VAL) often form hydrophobic or so-called (ILV)- cluster that prevent the intrusion of water molecules and serve as cores of stability in high-energy partially folded states (Kathuria *et al*., 2016). Although still not well understood, hydrophobic clusters seem to play a key role in protein stability (Basak *et al*., 2019). An available source to compute hydrophobic clusters is the BASIC web server (BASiC Networks), which relies on the CSU algorithm (Sobolev *et al*., 1999, 1996; Wołek *et al*., 2015). The CSU algorithm was released as a web server application but is unfortunately no longer maintained. We thus implemented the original CSU algorithm in Protlego to enable its use in a high-throughput fashion. The `compute_hydrophobic_clusters` function allows computing cluster for user-defined selections and visualizing them in the protein structure. **Fig. S1** summarizes the algorithm, which proceeds as follows: Two atoms A and B are considered to be in contact if a solvent molecule placed at the surface of A’s sphere, overlaps with the sphere formed by a solvent molecule plus the Van der Waals sphere of atom B (Sobolev and Edelman, 1995). The atoms are considered spheres of fixed radius, obtained from a previous publication (Shannon, 1976). If at any position a water molecule penetrates several atoms’ spheres, the contact is considered to belong to that whose centre is closest to the centre of atom A. In practical terms, Protlego defines an Atom class for each heavy atom that fulfils the user selection (such as residues ILE, VAL, and LEU).

During instantiation, the Atom class retrieves the coordinates of the neighbouring atoms. These are atoms that are closer than the sum of the two Van der Waals radii, each enlarged by the radius of the water molecule (1.4). Hence, for two carbon atoms to be considered candidates for atomic contacts they must be within 6.56 Å. The `Atom` class then discretizes the sphere of the atom in question into many uniform small sections. We use the Fibonacci grid (González, 2010; Wołek *et al*., 2015) to perform the discretization and select 610 points by default. The area corresponds to 0.0016 of the total area of the sphere. The algorithm then evaluates if any of the 610 (or a user-defined number) sections overlaps with the neighbours, and if so, the contact in the section is declared to belong to the sphere whose centre is closest to A’s centre.

The algorithm is followed for all the atoms until a matrix of residue-against-residue areas is computed. By default, we define that two residues are in contact when they have an overlapping area of at least 10 Å^2^. The adjacent matrix is converted to a graph, where every component corresponds to a (hydrophobic) cluster. The total area of the cluster is computed by the sum of the individual residue areas that comprise it.

## 3 Results

Here we present Protlego’s main applications by showcasing the example of the P-loop and Rossman α/β superfolds. The P-loop NTPases are a superfamily of enzymes that catalyse the hydrolysis of nucleoside triphosphate molecules (NTP). Despite extreme sequence and topology divergence, P-loop proteins are characterized by the presence of a sequence pattern GxxxxGKS/T (x being any residue) known as Walker A motif (Walker *et al*., 1982), which binds the terminal phosphate groups of NTPs. The P-loop is characterized by the Walker motif and the flanking β-strand and α-helix (Romero Romero *et al*., 2018). Due to their extreme sequence dissimilarity, several attempts have been made to classify P-loop proteins (Lupas and Martin, 2002; Pathak *et al*., 2014; Leipe *et al*., 2003). The SCOPe database classifies P-loop NTPases within the c.37 fold, all belonging to the c.37.1 superfamily (P-loop containing nucleoside triphosphate hydrolases). The domains are further divided into 27 families based on their β-sheet topologies. While all P-loop NTPases have three layers, with two and three helices sandwiching a 5-stranded parallel β-sheet, the order of the β-strands varies. For the great majority of P-loop kinases the order is 23145, as shown in **Fig. 3a**.

**Figure 3:**
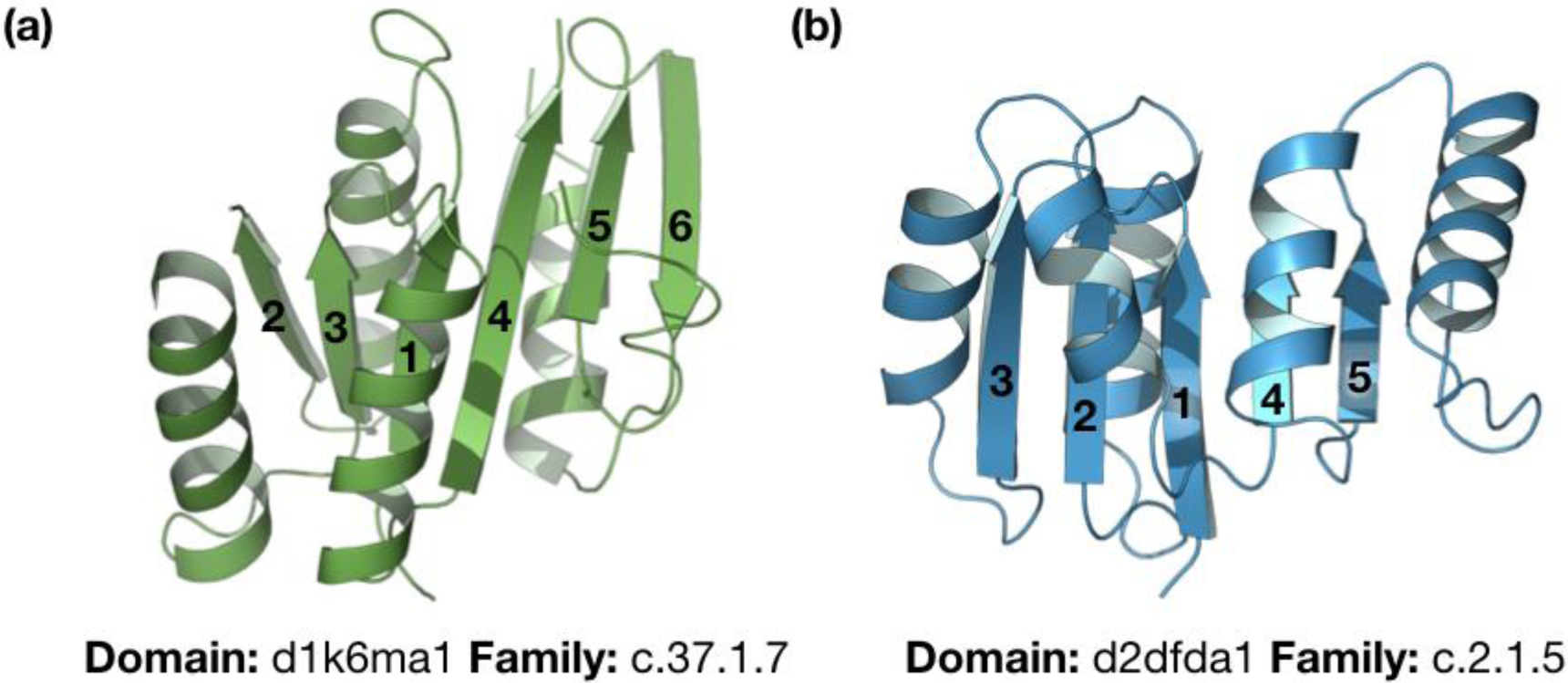
Exemplary topologies of the P-loop (a) and Rossman folds (b). Both are folds belonging to the α/β class. Whereas the P-loop containing nucleoside triphosphate hydrolases (SCOPe classification: c.37) have a 5- or 6-stranded β-sheet in the order 23145(6), the NAD(P)-binding Rossmann fold (SCOPe classification: c.2) has a 6-stranded β-sheet with an order of 321456.

The Rossmann fold is another of the most ancient and functionally diverse folds, catalysing more than 300 enzymatic reactions (Bukhari and Caetano-Anollés, 2013). Like the P-loop fold, most Rossmann enzymes use NTPs as cofactors and are formed by an α/β sandwich, in this case formed by 6 parallel β sheets with the order 321456 (**Fig. 3b)**. Similar to P-loop NTPases, Nicotinamide adenine dinucleotide (NAD)- and flavin adenine dinucleotide (FAD)-utilizing enzymes contain a Gly-rich motif that resides between α1 and β1, and are able to bind the phosphate groups of several NTPs. Besides, Rossman domains usually present an Asp/Glu residue at the top of β2 that provides a conserved and well-studied carboxylate-ribose bidentate interaction (Laurino *et al*., 2016).

The two folds comprise highly similar topologies, but deviate by the order of the core β-strands (23145 vs. 321456) and their different binding motifs in their N-termini. Furthermore, Laurino *et al*. studied that none of the P-loop NTPase families contain a canonical Rossman carboxylate-ribose interaction and that the nucleoside binding orientation in the two folds is reverse (Laurino *et al*., 2016).

Despite these differences, we explored the interaction between these α/β superfolds further, to (i) determine whether there are regions in their sequence where homology can be determined and (ii) showcase the application of the Protlego package on an interesting example. The code to reproduce these examples is provided in **Listing S1**.

### 3.1 Fetching fragments between the P-loop and the Rossmann folds

We first fetched all hits between the P-loop and Rossmann folds in the built-in Fuzzle database. The corresponding SCOPe identifiers for these folds are c.37 and c.2, respectively. We filtered for hits that present an HHsearch probability > 70%, an RMSD < 3 Å, fragment length between 10 and 200 amino acids, and TM-score below 0.3 in line with previous studies (Alva *et al*., 2015). The `Fetch_group` function retrieved 1737 hits for the two folds, containing 432 different unique domains. Each hit contains information about query and subject, start and end of the fragment they share, HHsearch probability, and RMSD, among others. The hits had an average length of 37.9 ± 17.0 amino acids (median 34.0) with a bimodal distribution with centres at 38 and 105 amino acids (**Fig. S2**), although markedly deviated towards short distances: 93% of the hits present a fragment length below 45 amino acids, and only 5% are over 100. Regarding other average properties, the hits have a mean RMSD of 2.3 ± 0.3 Å, an HHsearch probability of 77.0 ± 5 %, and a mean TM-score of 0.59 ± 0.1, which is indicative of a very good structural alignment (Xu and Zhang, 2010). Family composition does not cover all possible P-loop and Rossmann families. Out of the 26 families in the P-loop NTPase superfamily (c.37.1), 13 are not involved in these hits. From the side of the Rossman fold, out of 13 families in the NAD(P)-binding Rossmann-fold family (c.2.1), four do not show hits with P-loop domains above the cut-offs. **Figure S3** depicts the number of hits between each family pair: leaving apart hits between the automated matched families (c.37.1.0 and c.2.1.0), the majority of hits involve the c.37.1.10, c.37.1.1 or c.2.1.2 families.

The hits between P-loop and Rossmann domains reveal that these two folds appear to have homologous fragments, possibly at different regions of their sequences. In order to understand how many different types of fragments there are, their lengths, and specific location in the proteins’ sequences, we used similarity networks.

### 3.2 View of evolutionary relationships via networks

Similarity networks allow the study and visualization of the protein universe or subregions of it, where the nodes represent protein domains and the links connect two domains when they have a fragment in common. In this case, we fetched 1737 hits which overall contain 432 different domains. All domains in the Fuzzle database have been clustered previously according to the regions in their sequence (Ferruz *et al*., 2020). For example, a domain A might have two hits: in hit1 it forms a local alignment with amino acids 1 to 30, while in hit2 it aligns with amino acids 90 to 150. Because these two areas are virtually different fragments, we denote these regions as two different clusters. In this case, the 432 unique domains in our selection belong to 460 clusters.

The similarity network for these hits is shown in **Fig. 4**. Domains belonging to the P-loop and Rossmann folds are coloured in green and blue, respectively. The network contains 460 nodes and 1213 links. Each of the ‘island-like’ motifs correspond to a set of domains that contain a common fragment. These ‘island-like’ motifs are termed *components* in network theory. For further details on the construction of the network and definitions we refer to our previous publication on Fuzzle (Ferruz *et al*., 2020). The network consists in this case of 17 components with very diverse sizes. The largest component is composed of 361 nodes, whereas 7 components present only 2 nodes. This is a known behaviour in network theory applied to biological systems: while a few nodes have many connections, most nodes are only sparsely linked to others (Barabási and Albert, 1999; Wuchty, 2001; Levitt, 2009). We will focus in this analysis on the four most populated components. A summary of their properties is presented in **Table 2**.

**Table 2:**
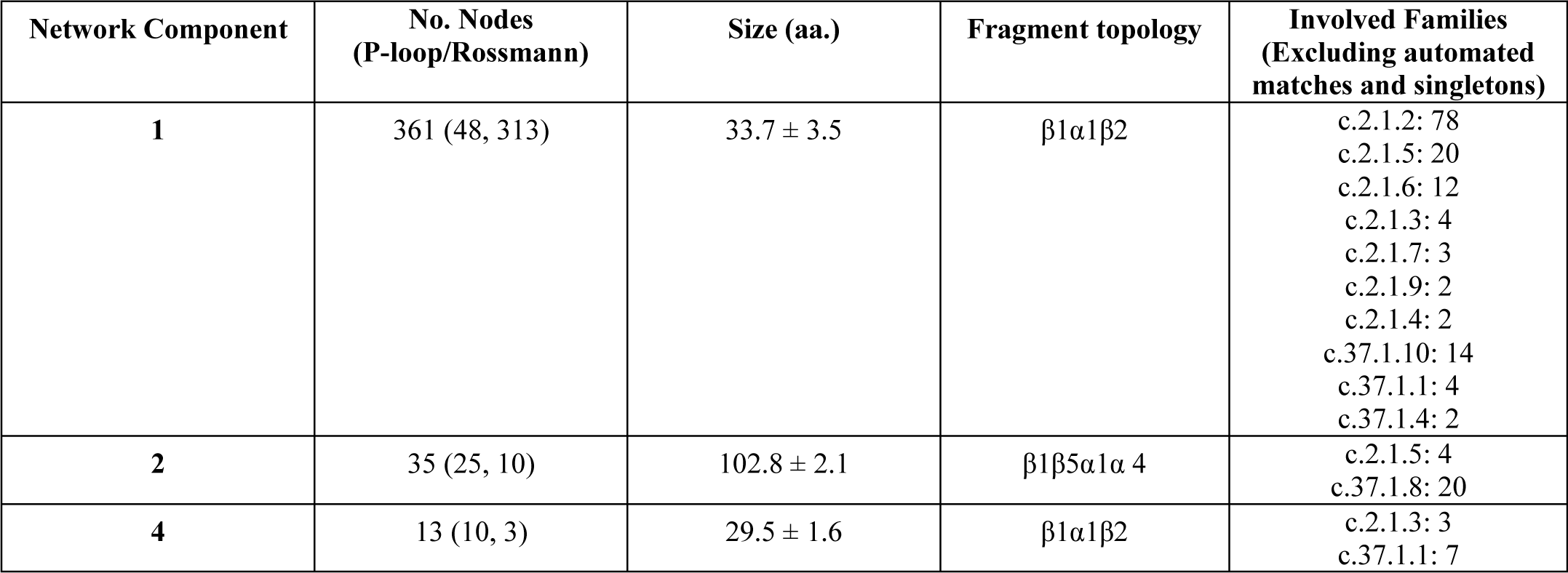
Summary of properties for components 1, 2, and 4 in the similarity network.

| Network Component | No. Nodes<br>(P-loop/Rossmann) | Size (aa.) | Fragment topology | Involved Families<br>(Excluding automated matches and singletons) |
| --- | --- | --- | --- | --- |
| 1 | 361 (48, 313) | 33.7 ± 3.5 | $\beta 1\alpha 1\beta 2$ | c.2.1.2: 78<br>c.2.1.5: 20<br>c.2.1.6: 12<br>c.2.1.3: 4<br>c.2.1.7: 3<br>c.2.1.9: 2<br>c.2.1.4: 2<br>c.37.1.10: 14<br>c.37.1.1: 4<br>c.37.1.4: 2 |
| 2 | 35 (25, 10) | 102.8 ± 2.1 | $\beta 1\beta 5\alpha 1\alpha 4$ | c.2.1.5: 4<br>c.37.1.8: 20 |
| 4 | 13 (10, 3) | 29.5 ± 1.6 | $\beta 1\alpha 1\beta 2$ | c.2.1.3: 3<br>c.37.1.1: 7 |

**Figure 4:**
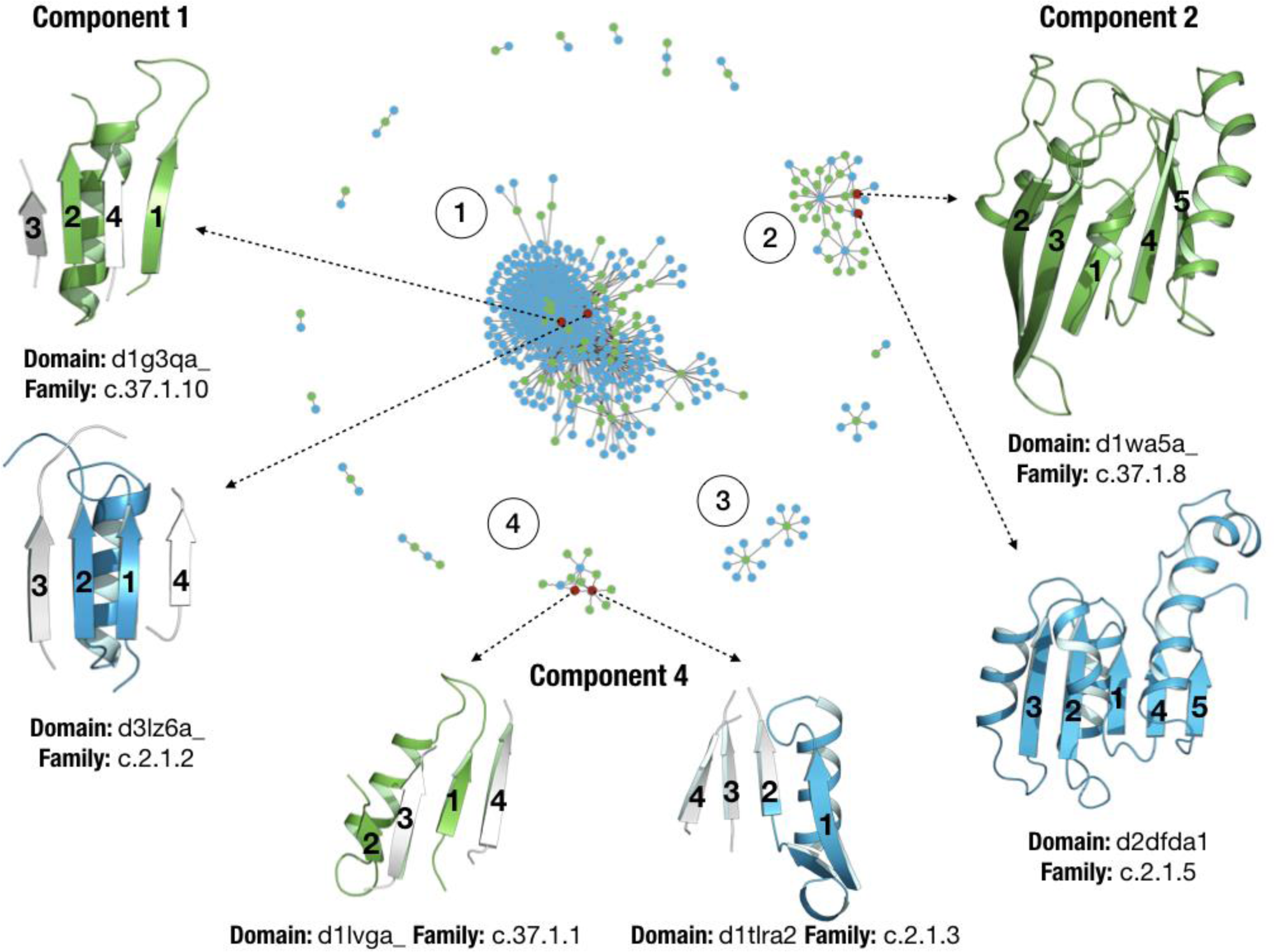
Similarity network for hits between the P-loop and Rossmann folds that surpass the thresholds (see main text). Nodes coloured in green and blue represent P-loop and Rossmann domains, respectively. Representative domains and their fragments in common are shown for components 1 (a), 2 (b) and 3 (c).

With 361 nodes component 1 is the major component of the network. The populations somewhat deviate towards Rossmann domains, with 48 and 313 nodes for the P-loop and Rossmann folds, respectively. These numbers can be appreciated in the component topology, with most Rossmann connecting around P-loop nodes that act as *hubs*. Not surprisingly, the 16 most connected nodes in the component correspond to P-loop domains. Interestingly, all these nodes belong to either the family c.37.1.10 (Nitrogenase iron protein-like) or the c.37.1.0 family (automated matches). The Nitrogenase iron protein-like family (NifH) is involved in the fixation of nitrogen and has seven β-sheets in the order 3241567. Most of the Rossmann domains on the other hand belong to the c.2.1.2 family (tyrosine-dependent oxidoreductases). The nodes in this component share a fragment with an average length of 33.7 ± 3.5 amino acids. The fragment is located at the N-terminal end in both folds and consists of the β1α1β2 fragment, which contains the Walker and Gly-rich motif. **Fig. 3** shows two representative P-loop and Rossmann domains, d1g3qa_, the cell division regulator MinD from the c.37.1.10 family (Hayashi *et al*., 2001) and d3lz6a, 11-beta-hydroxysteroid dehydrogenase 1 protein from the c.2.1.2 family (Cheng *et al*., 2010). Representative domains shown in **Fig. 3** for each component are also depicted in red in the network.

The second largest component (no. 2) contains 35 nodes, with 25 and 10 P-loop and Rossmann domains, respectively. The fragment these domains have in common has an average length of 102.8 ± 2.2 amino acids and comprises a fragment defined by the first five sheets and four helices, practically spanning the entire domain structure (**Fig. 3)**. The most connected domain is d2dfda1, a malate dehydrogenase NAD binding Rossmann domain (Ugochukwu *et al*.), with 18 connections. **Fig. 3** shows two representative domains, d1wa5a_, a G-protein with an antiparallel β2, and d2dfda itself. The domains in this component, besides the families of automated matches, belong to the families c.37.1.8 (G-proteins) and c.2.1.5 (LDH N-terminal domain-like).

Component 3 corresponds to a subgraph with 15 nodes, with 2 P-loop domains that connect to 13 Rossmann nodes, each connecting 7 domains. The Rossmann domain d2yv1a1, a succinyl-CoA binding domain (Niwa *et al*.), acts as a bridge between the two P-loop domains (**Fig. 3c**). They correspond in fact to the same domain d2g0ta1, that due to slight differences in the sequence where the fragments are present got assigned to different clusters (d2g0ta1_2 and d2g0ta1_13). This domain belongs as well to a domain from the Nitrogenase iron protein-like family (NifH), but when looking in detail at its structure, we observe that it contains an extra N-terminal Rossmann fold which has not been classified by SCOPe as a different domain. The hit with the Rossmann domain d2yv1a1 is thus expected as both domains have the same fold. Component 3 exemplifies a case of misclassification occurring sometimes in hierarchical databases and as such it is not presented in detail in **Table 2** and **Fig. 3**.

Lastly, component 4 contains 13 nodes, with 3 Rossmann domains connecting to 10 P-loop domains. The most connected domain is d1t1ra2, a 1-deoxy-D-xylulose-5-phosphate reductoisomerase belonging to family c.2.1.3, with 7 connections (Yajima *et al*., 2004). The fragment in this component is short, with an average length of 29.5 ± 1.6 amin acids. It also corresponds to the N-terminal βaβ fragment containing the conserved motifs. Major differences with component 1 are the slightly smaller size of this fragment and the different families which contain it. While component 1 mainly contains c.37.1.10 domains, component 4 contains domains of the nucleotide and nucleoside kinase family (c.37.1.1). In this case the network has discerned fragments into different components that are present in different P-loop families. **Fig. 3** depicts this fragment in two representative domains, d1lvga_ (Sekulic *et al*., 2002), a Guanylate kinase from the nucleosides kinases family (c.37.1.1) and d1t1ra2 from family c.2.1.3, previously mentioned in component 3. Interestingly, the fragment in the Rossmann domains corresponds to βaβ where the second beta sheet is a short insertion before β2, which corresponds in the P-loop structures to an exceptionally short β2 strand.

### 3.3 Automatic construction of chimeras

Protlego builds all possible chimeras between two domains based on their sequence alignment (**Methods, Fig. 2**). Here we have created all possible chimeras between the all P-loop/Rossmann pairs of domains. As we had previously noticed a misclassification of domain d2g0ta1, which contains an extra N-terminal Rossmann domain, we removed all hits containing this domain, which led to 1693 hits (previously 1731). We first performed chimeragenesis using a global alignment (**Methods**), which lead to a total of 1158 chimeras. Remarkably, only 5% of hits led to chimeras without backbone clashes, with hits presenting short alignment lengths producing very few chimeras (**Fig. S5a**). Besides the average short length of these hits and its subsequent lower number of possible fusion points, another reason for the low number of chimeras is the intrinsic topology of the β strands in the parents. The shifting of β2 in the β-strand order (23145 vs. 213456) leads to backbone clashes once the two termini are combined. We decided to perform partial alignments instead, to see if stepwise alignments of shorter motifs improved the statistics. Indeed, the partial alignment provided 2503 chimeras coming from 27% of hits, with hits with shorter lengths producing more chimeras than before (**Fig. S5b**). We then had a look at those combinations of families that produced most chimeras. The combination of families c.37.1.8 - c.2.1.5 led to 709 chimeras, despite only having 34 hits (**Fig. S6**). In second place, the pair between families c.37.1.1 and c.2.1.3, presented 17 hits and produced a total of 104 chimeras. The third most abundant pair corresponds to families c.37.1.10 and c.2.1.2, which with 202 hits gave rise to 48 chimeras. Remarkably, these three examples exactly correspond with the predominant families found in component 2, 4, and 1, respectively (**Fig. 4**).

We decided to explore the first example in detail. Concretely, we built all possible chimeras between domains d1wa5a_ and d2dfda1, already chosen as representatives of component 2 in **Fig. 4**. The hit has a HHsearch probability of 81.7%, a fragment length of 101 amino acids that superimpose with an RMSD of 2.89 Å over 85 Cα atoms and a TM-score of 0.55. The alignment identity is 14%. The algorithm first maps all amino acids in the sequence alignment to their corresponding positions in the PDB (**Fig. 2**). In this case, the 101 amino acids were successfully mapped to the structures. Then, the partial alignment divided the (βα)1-4β5 fragment into six sections and distances between Cα pairs were computed. From the 101 aligned positions, a maximum of 202 offspring chimeras could be expected in the ideal scenario when the structures align perfectly, and the resulting chimeras do not have backbone clashes. This scenario is rather unrealistic, as it would require an RMSD of 0 Å in proteins with different topologies. In this case, only 32 of those 101 points presented a distance between Cα atoms below the default cut-off of 1 Å. From the possible 64 chimeras altogether in combination1 and 2, 43 of the built chimeras did not pass the last quality filter due to backbone clashes. Overall, 21 chimeras passed all the criteria and were successfully built. **Fig. S7** summarizes the outcome of each fusion point. An equivalent plot can be obtained by calling the Builder.plot_chimeras() function.

**Fig. S8** represents the 21 chimeras coloured according to the fragments they inherited from their parents (d1wa5a_: P-loop, green, d2dfda1: Rossmann, blue). Chimeras in combination1, (with d2dfda1 at the N-terminus) have the topology 321456, whereas the topology for chimeras in combination2 is strand order 23145.

### 3.4 Energetic evaluation

We energetically evaluated the 21 chimeras with the Amber forcefield both with and without backbone flexibility enabled. Results are summarized in **Table 3**. Chimera scoring could be useful in contexts when only a few chimeras can proceed to the experimental assessment. In this case, fast means to rank them computationally are necessary. As expected, the chimeras tend to score better when allowing backbone rearrangements during minimization. When ranking the chimeras by their score per residue, we observe that the first four correspond to the chimeras in combination2 with the most P-loop content (comb2_109-118). In the middle part of the table we find mostly chimeras from combination1 that contain 3 β-strands from each parent. The last part of the ranking is mostly populated with combination2 chimeras with less P-loop content (comb2_80-107). One possibility explaining the striking differences observed between chimeras from comb2_80-107 and chimeras comb2_109-118 is that the first group involves a fusion point right in the middle of the helix coming from β4 while the four chimeras in comb2_109-118 conserve the native helix from the P-loop domain (**Fig. S8**). We have also minimized and scored the parent domains to allow direct comparison. Interestingly, the parent domains have scores at the two extremes of the chimera distribution: domain d1wa5a_, with a potential energy per residue of −22.1 kcal/mol confers the better scoring domain. On the other side of the spectrum, domain d2dfda1 presents a potential energy of −18.1 kcal/mol, positioning itself at the bottom of the table. Given these results, we questioned whether the order in Table 3 reflects the partial content of the two chimeras, with higher-scoring chimeras having more P-loop content. Although some trend is observed for the first chimeras, with high P-loop content and very negative scores, an overall trend between the two variables could not be found (R^2^ =0.25). These findings were perhaps to be expected, as potential energy calculations depend on a myriad of factors and subtle changes in protein packing may dramatically affect their outcome.

**Table 3:**
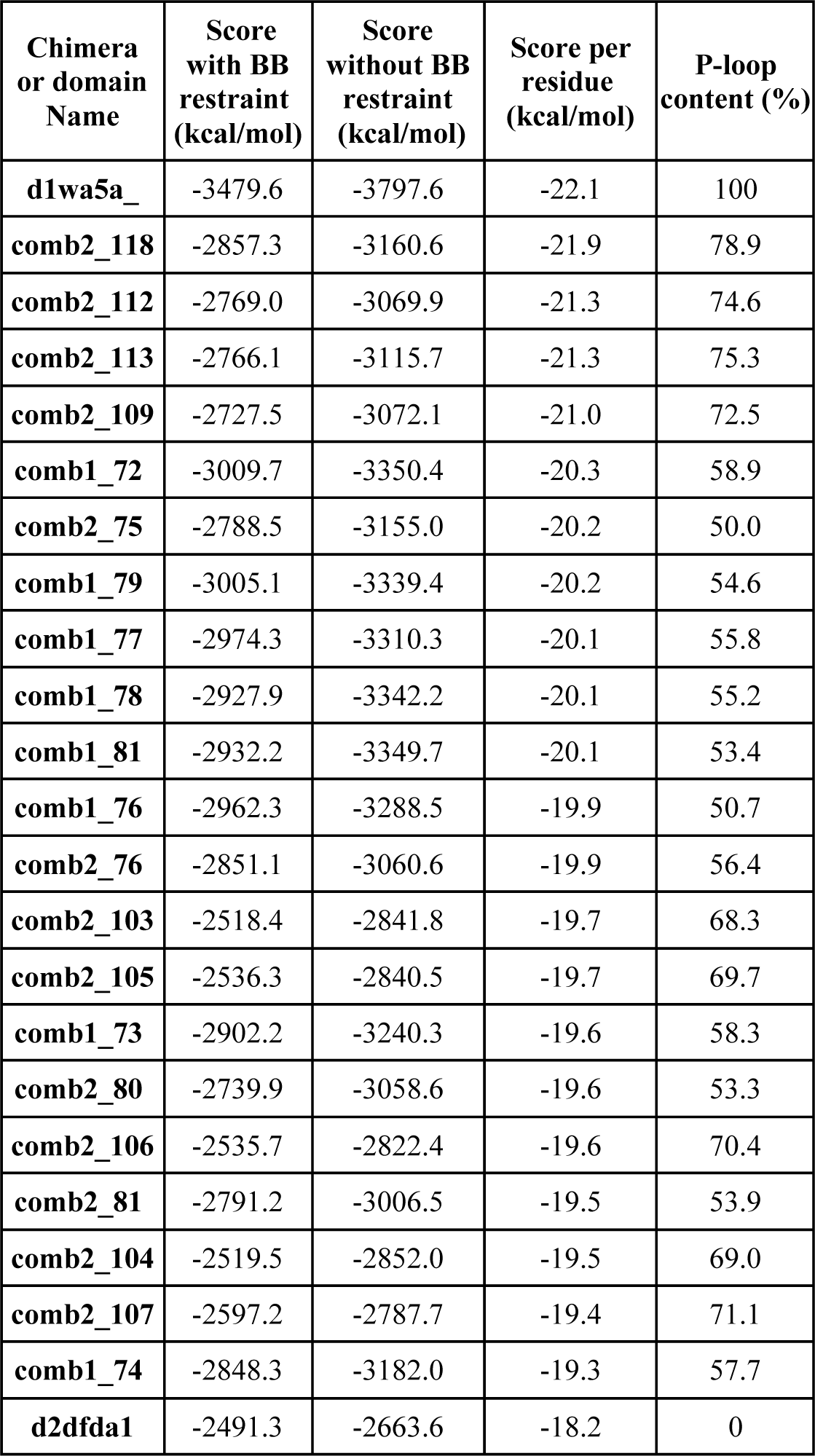
Summary of the energy minimization with the Amber forcefield. The minimizations were performed allowing for backbone relax or not. The table is ordered according to the best scoring chimeras without backbone restraints.

### 3.5 Structural analysis

Protlego enables the analysis of several structural features, such as hydrophobic clusters, hydrogen bond plots (Bikadi *et al*., 2007), salt-bridge and hydrogen networks, solvent-accessible surface area (SASA) calculations, contact orders, and contact map representations, among others. Here, we showcase some of these analyses with chimera comb1_72, as it provides an interesting structure with three inherited β-strands from each parent and as it is one of the highest scoring members in the set.

We started by analysing hydrophobic clusters (**Methods**). To allow direct comparison, we performed the same computation on its parents d1wa5a_ and d2dfda1, in all cases after a non-restraining minimisation. The two parents contain two main hydrophobic clusters, flanking on both sides of the β-sheet (**Fig. S9**). While clusters in domain d2dfda consist of 16 (black) and 15 (white) residues, both clusters in domain d1wa5a_ are composed of 9 residues, despite d1wa5a_ containing more residues overall (172 vs. 147). We have often observed that the chimeras do not inherit the hydrophobic cluster conformation from its parent proteins, due to non-hydrophobic new residues in places that can break the cluster continuity. However, comb1_72 contains two major clusters consisting of 16 and 11 residues (**Fig. 5c and Fig. S9c**). Although the largest cluster fails to reproduce the large area of the Rossmann parent d2dfda1 (1996.7 vs 1755.3 Å^2^) it contains residues from the parent P-loop domain d1wa5a_, forming a continuous entity. A similar behaviour is observed for the second cluster which presents an area of 1519.3 Å^2^ and with 11 residues spans the two regions.

**Figure 5:**
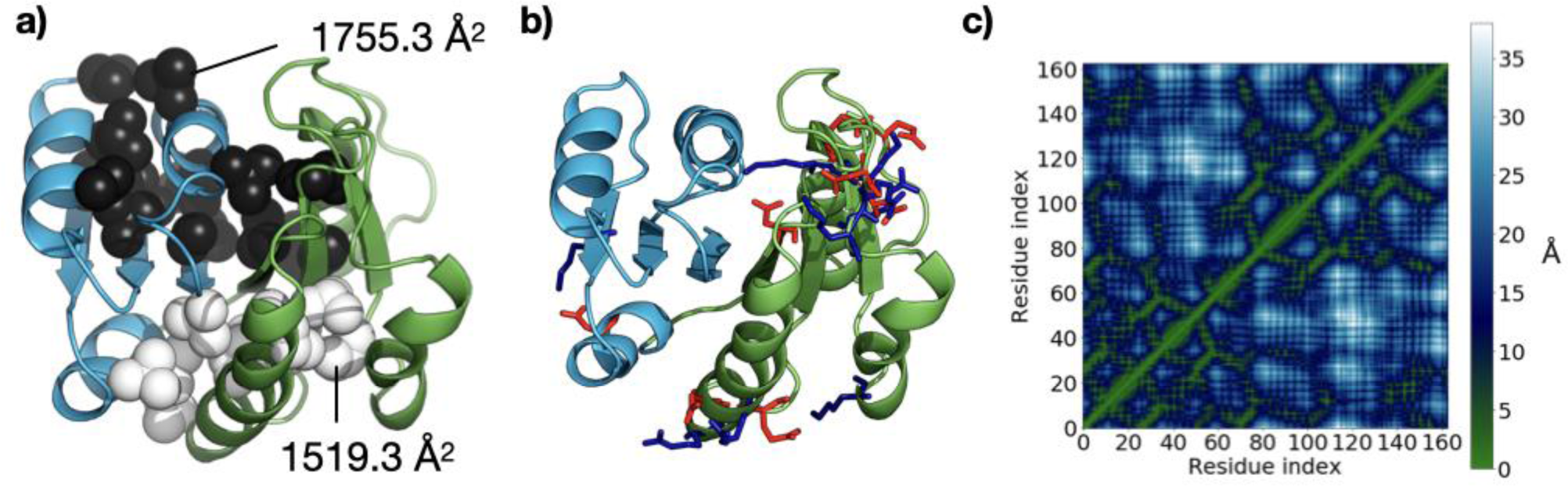
Some of the structural analysis included in Protlego. **a)** The two largest hydrophobic clusters shown in black and in white computed for chimera comb1_72. **b)** Computation of salt bridges. Acidic residues are shown in red and basic ones in blue. c) Computation of contact maps. Distances color-coded and listed in Å: Regions in green are close in distance whereas those in blue are far apart.

We then turned our interest to the computation of salt bridges. The Rossmann domain contains fewer salt bridges, summing up to only five (**Fig. S10a**). The P-loop domain, however, has a total of 14 salt-bridges, mostly located in the region defined by the last three β-strands (**Fig. S10b**). Comb1_72 contains 9 salt bridges, 8 of them inherited from parent d1wa5a_ (**Fig. S10c**). Comb1_72 forms a new interaction between β3 and α3, perhaps a consequence of repacking between α3 and α4 after minimisation. The salt-bridge between β1 and β4 on the other hand has been depleted after fusion.

On a different note, contact maps reduce the dimensionality of proteins to a 2D-plot that reveals what positions in the protein are close in distance. An advantage of contact maps is that they are invariant to rotations and translations and are often used for protein superimposition (Holm and Sander, 1996). As expected, the chimera contact map presents a combination of the contact maps of their parents, with residues 1-71 reflecting the Rossman map, and residues 71 – 165 resembling the P-loop map (**Fig. 5** and **Fig. S11**).

Another interesting property is the contact order, which measures the average sequence distance between residues that are in structural contact. Contact orders have been studied in the context of protein folding, and it is suggested that higher contact orders indicate longer folding times (Plaxco *et al*., 1998). Interestingly, designed proteins tend to have lower contact orders than the distribution observed in natural proteins (Bonneau *et al*., 2002). Their typical contact order value ranges anywhere from 5 to 25%. In line with these values, parent domains and chimera have a contact order of 14.1 % (d2dfda1), 18.3 % (d1wa5a_), and 15.2 % (comb1_72), respectively. Solvent-accessible surface area is the area of a protein that is accessible to a solvent molecule. In Protlego, it is also possible to compute the specific contribution of polar and apolar residues to the total surface. For chimera comb1_72 we calculate a total area of 84.7 nm^2^, with a contribution from polar and apolar residues of 55.9 nm^2^ and 27.4 nm^2^, respectively. The Rossmann domain d2dfda1 has a total area of 73.4 nm^2^ with contributions of 40.6 nm^2^ (polar) and 31.9 nm^2^ (apolar). Lastly, the P-loop domain d1wa5a_ contains surfaces of 88.7 nm^2^ in total, with 60.6 nm^2^ from polar and 27.0 nm^2^ from apolar residues.

## 4 Conclusion

The design of novel proteins via recombination of sub-domain sized fragments provides an attractive new route for the design of customizable proteins. The modelling and ranking of several hundreds of chimeric proteins prior to its testing in the lab requires however the use of several techniques and a considerable degree of expertise. Here, we have implemented Protlego, a python-based open-source software for the automatic construction of chimeras and its structural analysis. Protlego is divided into five applications. First, it contains a lightweight version of the Fuzzle database which enables fetching hits from specific SCOPe groups or that fulfil a specific user-defined criterion. The retrieved hits can then be represented via similarity networks, facilitating the understanding of convoluted evolutionary relationships. Selected hits from Fuzzle can then be used to build chimeras with one recombination point, whose structural features can be further analysed in detail, or its potential energies estimated and ranked.

In this work, we chose to showcase Protlego’s features on a biologically relevant example which has attracted the attention of many groups in the past: The relationship of the P-loop and Rossmann folds. By fetching hits from the database, we obtained a total of 1693 domains of these two folds. We represented these hits via a network, which nicely separated their different fragments. For example, the fragment in component 1 and 4 is a rather short motif of ∼30 amino acids, located at the N-terminus of the sequence, whereas the fragment in component 2 is a large fragment spanning 5 β-strands. Interestingly, the network separated also the fragments by family relationships, with each component preferably containing a family pair. In fact, not all P-loop and Rossmann families appear homologous, rather only a subset of them, with families c.37.1.10 and c.2.1.2 being predominantly connected (**Fig. 4, Fig. S5**). We selected the hit between domains d1wa5a_ and d2dfda1 to illustrate the process of automatic chimera construction. In particular, this hit leads to 21 chimeras with different parent contents and estimated potential energies (**Table 3**). We selected one chimera for further analysis, which showed that it could be potentially well folded. Specifically, the hydrophobic clusters coming from the two parents coalesced nicely spanning the two combined regions, and other features showed values similar to those of natural proteins.

The code for all the analysis in this work is provided. We believe that Protlego can be a useful tool for the evolutionary biologist and protein engineer alike. Furthermore, it is our goal to allow contribution from the scientific community by providing full access to the code repository, which can be added in future Protlego releases. Examples, documentation and code are freely available at https://hoecker-lab.github.io/protlego/

## Acknowledgements

We thank Francisco Lobos for discussions on the code.

## Funding

This work has been supported by the European Research Council (ERC Consolidator grant 647548 ‘Protein Lego’) and the Volkswagenstiftung (grant 94747).

## Conflict of Interest

none declared.

## Notes

https://hoecker-lab.github.io/protlego/

### Competing Interest Statement

The authors have declared no competing interest.

