## Supplementary Information for "Protlego: A Python package for the analysis and design of chimeric proteins"

*Supporting Information for:*

### Protlego: A Python package for the automated design of protein chimeras and structural analysis.

### LISTINGS

#### Listing S1: Code used to generate the results in the manuscript.

```
# Importing the module
from protlego import *

# Retrieving all hits between the two folds
hits=fetch_group('c.37','c.2')

# Creating network and plotting
a=Network(hits)
graph = a.create_network()
a.plot_graph(graph,'fold')

# Building all possible chimeras.
chimeras={}
for index,hit in enumerate(hits.hits):
    a=Builder(hit)
    aln=a.get_alignment(hit.query,hit.no)
    a.superimpose_structures(aln,partial_alignment=True)
    chimeras[index]=a.build_chimeras(partial_alignment=True)

# Selecting one hit and scoring its chimeras
selected_hit=hits[1069]
b=Builder(selected_hit)
aln=a.get_alignment(selected_hit.query, selected_hit.no)
b.superimpose_structures(aln,partial_alignment=True)
sel_chimeras=b.build_chimeras(partial_alignment=True)
b.plot_curves()
values_amber={}
chimeras_after_amber={}
chimeras_after_amber={}
for key,chimera in chimeras.items():
    values_amber[key],chimeras_after_amber[key]=minimize_potential_energy(chimera,
'amber', cuda=True, restraint_backbone=False)

# Structural analysis of chimera comb1_72
chimera = chimeras_after_amber['comb1_72']
clusters = chimera.compute_hydrophobic_clusters()
chimera.view()
chimera_salts = chimera.compute_salt_bridges()
chimera.view()
calc_sasa(chimera)
calc_contact_order(chimera)
distance_matrix=calc_dist_matrix(chimera,type='distances', plot=True)
hnets=chimera.compute_hydrogen_networks()
calc_contact_order(chimera)
chimera.view()
```

### FIGURES

**Figure S1: Summary of the algorithm to compute hydrophobic clusters.** Given a PDB, an Atom class is defined for each heavy atom in the given selection (by default ILE, VAL and LEU residues) (**1**). The atom is represented as a sphere whose surface is divided into 610 sections (or a user defined number). Each section is evaluated whether it intersects another Atom object or not (**2**). The total areas are summed up per residue and a matrix is built (**3**). The matrix can be transformed into a graph (**4**), whose components virtually correspond to the fragments in the protein (**5**)

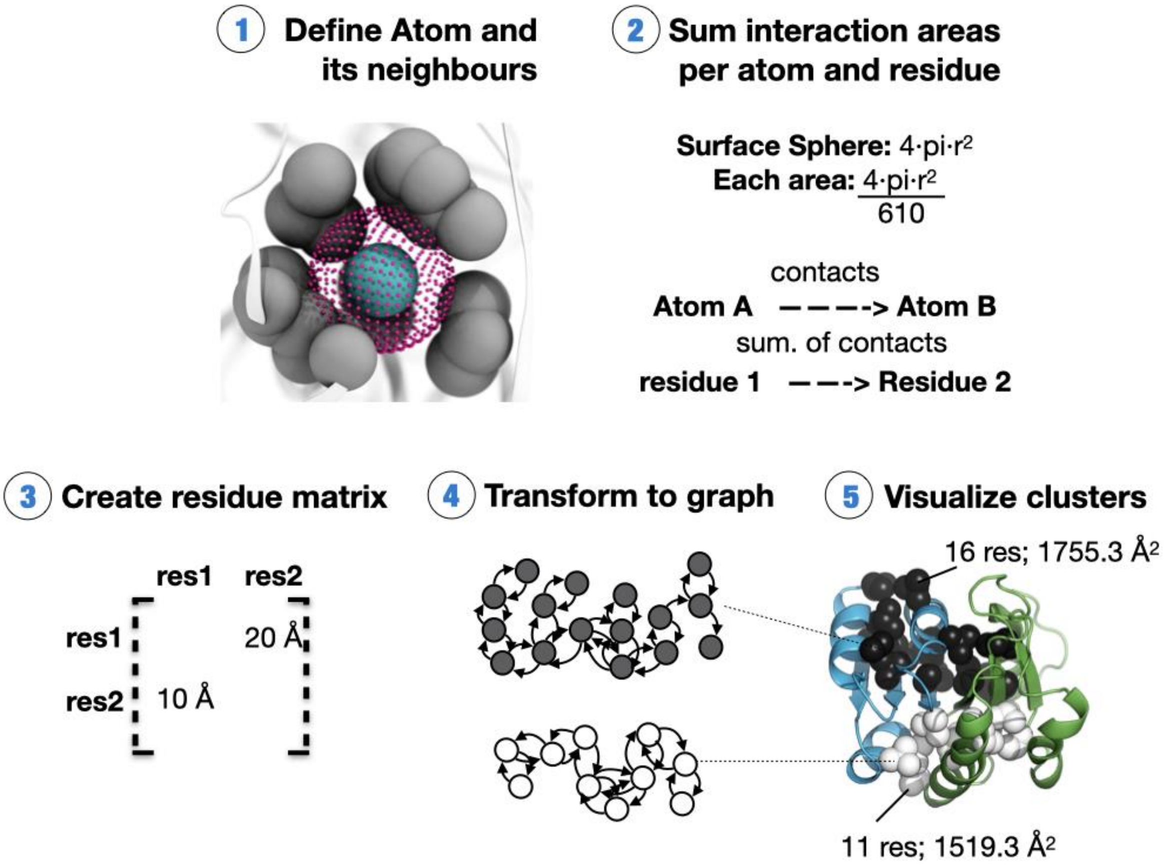

**Figure S2: Distribution of fragment length for fragments in the retrieved hits.** In blue, the histogram including all hits (1737), in orange, excluding domain d2g0ta1 due to misclassification (1693). The two sets have virtually identical distributions.

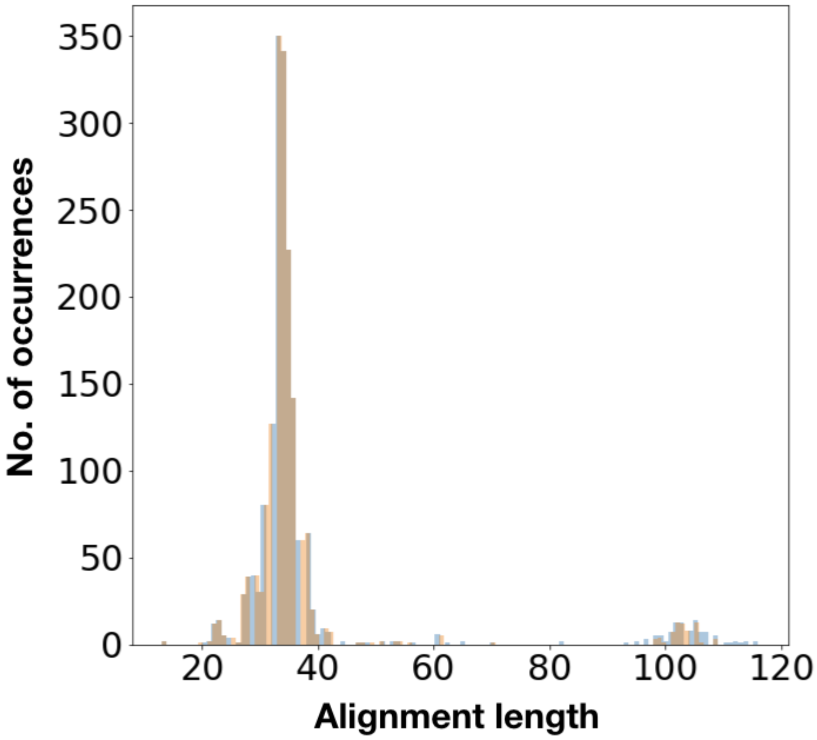

**Figure S3: Number of hits found between each P-loop and Rossmann family.** Besides the hits between automated matches (c.37.1.0 and c.2.1.0), the majority of hits involve the c.37.1.10 or c.2.1.2 families. Column and row 1 are depicted in gray as they represent the families of automated matches. Column 10' indicates the number of chimeras produced by family c.37.1.10 when removing hits that include domain d2g0ta1 as query or subject due to misclassification (see main manuscript). Greater numbers are depicted in darker shades of green.

P-loop families 'c.37.1.x' (columns); Rossmann families 'c.2.1.x' (rows)

| | 0 | 1 | 2 | 3 | 4 | 5 | 6 | 7 | 8 | 9 | 10 | 10' | 11 | 12 | 13 | 14 | 15 | 16 | 17 | 18 | 19 | 20 | 21 | 22 | 23 | 24 | 25 | 26 |
|---|---|---|---|---|---|---|---|---|---|---|---|---|---|---|---|---|---|---|---|---|---|---|---|---|---|---|---|---|
| **0** | 361 | 31 | | 2 | 6 | 2 | | | 23 | | 552 | 550 | | | | | | | | | 5 | | | | | | 1 | |
| **1** | 1 | | | | | | | | | | 1 | 1 | | | | | | | | | | | | | | | | |
| **2** | 177 | 14 | | 4 | 14 | | | | | | 202 | 202 | | | | | 4 | | | | | | | | | | | |
| **3** | 17 | 17 | | | | | | | | | 25 | 2 | | | | | | | | | | | | 2 | | | | |
| **4** | 7 | | | | | | | | | | 10 | 10 | | | | | | | | | | | | | | | | |
| **5** | 33 | 7 | | | | | | | 34 | | 37 | 37 | | | | | | | | | | | | | | | | |
| **6** | 37 | | | | | | | | | | 51 | 51 | | | | | | | | | | | | | | | | |
| **7** | 6 | | | | | | | | 1 | | 5 | 5 | | | | 2 | | | | | | | | | | | | |
| **8** | | | | | | | | | | | 19 | 0 | | | | | | | | | | | | | | | | |
| **9** | 10 | 2 | | | | 1 | | | | | 14 | 14 | | | | | | | | | | | | | | | | |
| **10** | | | | | | | | | | | | 0 | | | | | | | | | | | | | | | | |
| **11** | | | | | | | | | | | | 0 | | | | | | | | | | | | | | | | |
| **12** | | | | | | | | | | | | 0 | | | | | | | | | | | | | | | | |
| **13** | | | | | | | | | | | | 0 | | | | | | | | | | | | | | | | |

**Figure S4: Similarity network colored by families.** Each family in the network is automatically assigned a color by Protlego. The different components in the network have different family contents.

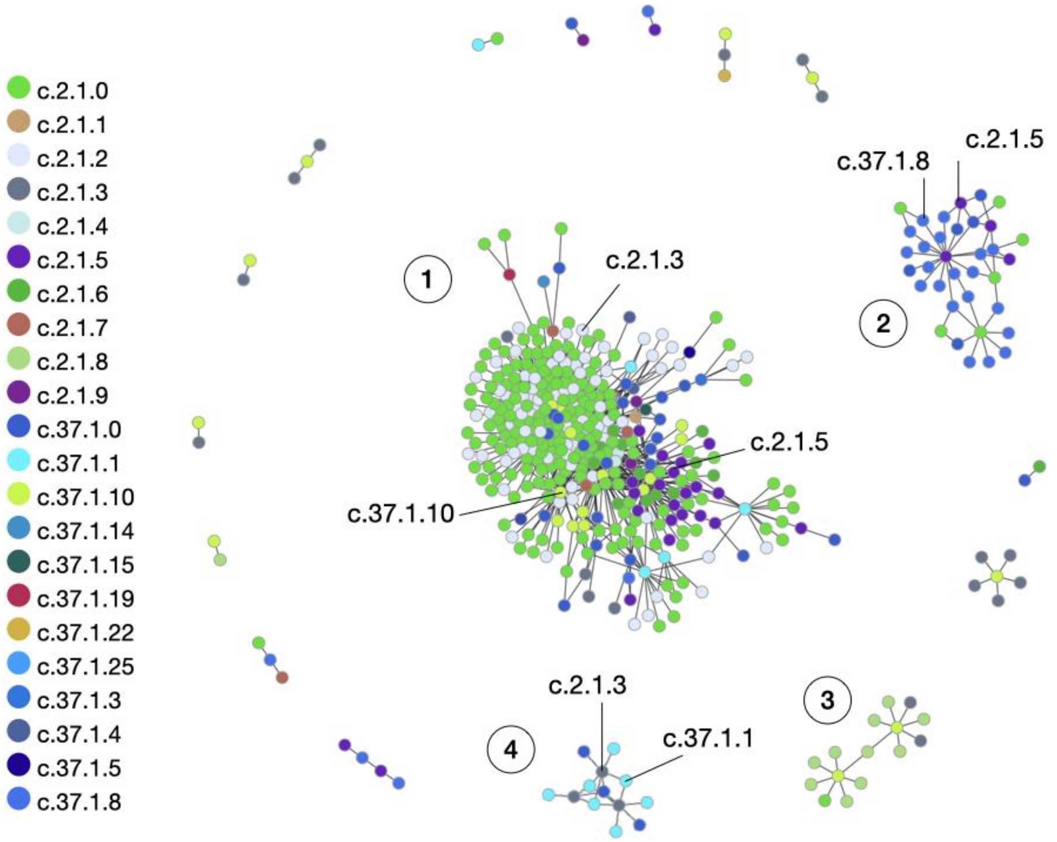

**Figure S5: Relationship of alignment length and number of produced chimeras for the global (a) and partial (b) alignments.**

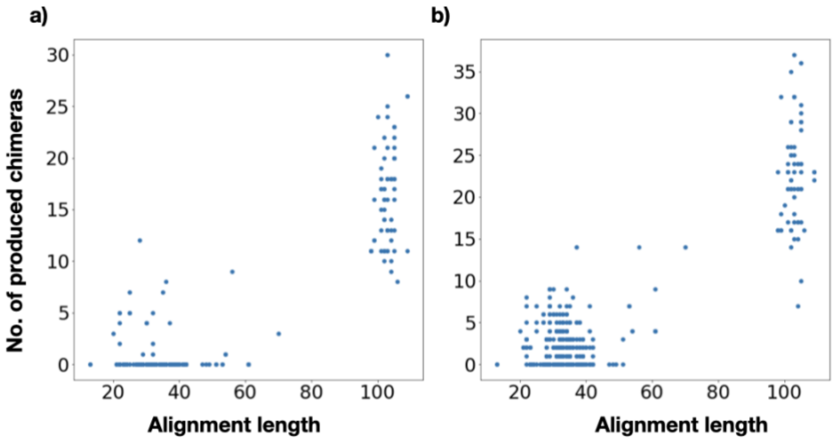

**Figure S6: Combination of families and the number of chimeras they produced.** The numbers correspond to the partial alignment algorithm. The total number of chimeras is 3170 when counting all domains, and 2503 when removing domain d2g0ta1. Column and row 1 are depicted in gray as they represent the families of automated matches. Column 10' indicates the number of chimeras produced by family c.37.1.10 when removing hits containing d2g0ta1 due to misclassification. The Figure is represented such as **Fig. S3**, with larger numbers being depicted in darker shades of green.

P-loop families 'c.37.1.x' (columns); Rossmann families 'c.2.1.x' (rows)

| | 0 | 1 | 2 | 3 | 4 | 5 | 6 | 7 | 8 | 9 | 10 | 10' | 11 | 12 | 13 | 14 | 15 | 16 | 17 | 18 | 19 | 20 | 21 | 22 | 23 | 24 | 25 | 26 |
|---|---|---|---|---|---|---|---|---|---|---|---|---|---|---|---|---|---|---|---|---|---|---|---|---|---|---|---|---|
| 0 | 329 | 71 | | 9 | | | | | 508 | | 206 | 188 | | | | | | | | | 25 | | | | | | | |
| 1 | 7 | | | | | | | | | | 4 | 4 | | | | | | | | | | | | | | | | |
| 2 | 112 | 16 | | 11 | 4 | | | | | | 48 | 48 | | | | | | | | | | | | | | | | |
| 3 | 54 | 104 | | | | | | | | | 152 | 0 | | | | | | | | | | | | 8 | | | | |
| 4 | | | | | | | | | | | 9 | 9 | | | | | | | | | | | | | | | | |
| 5 | 174 | 2 | | | | | | | 709 | | 32 | 32 | | | | | | | | | | | | | | | | |
| 6 | 35 | | | | | | | | | | | 0 | | | | | | | | | | | | | | | | |
| 7 | 3 | | | | | | | | | | 2 | 2 | | | | 14 | | | | | | | | | | | | |
| 8 | | | | | | | | | | | 497 | 0 | | | | | | | | | | | | | | | | |
| 9 | | 4 | | | | | | | | | 3 | 3 | | | | | | | | | | | | | | | | |
| 10 | | | | | | | | | | | | 0 | | | | | | | | | | | | | | | | |
| 11 | | | | | | | | | | | | 0 | | | | | | | | | | | | | | | | |
| 12 | | | | | | | | | | | | 0 | | | | | | | | | | | | | | | | |
| 13 | | | | | | | | | | | | 0 | | | | | | | | | | | | | | | | |

**Figure S7: Summary of chimera outcomes for each alignment position.** Out of the 101 alignment positions, 31 present distances between Cα pairs below 1 Å. 24 and 19 of these points produce a chimera with clashes (shown in black). 21 final chimeras are buildable for combination 1 (yellow) and 2 (red)

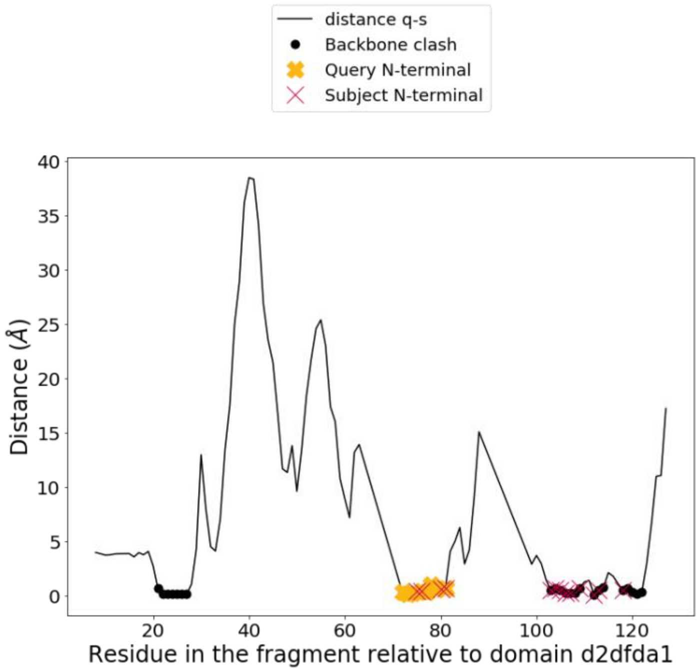

**Figure S8: Representation of all chimeras produced for the selected hit.** The hit between the domains d2dfda1 (query, Rossman) and d1wa5a_ (subject, P-loop) produced 21 offspring chimeras. Names for each chimera are depicted below its representation. The number summarizes the combination that it comes from (comb1 or comb2) and the residue where the parents are joined. Chimeras in combination 1 have a topology of strand order 321456, whereas chimeras in combination 2 the strand order 23145. The colouring method preserves the previous representation for the parents (blue: Rossmann, green: P-loop)

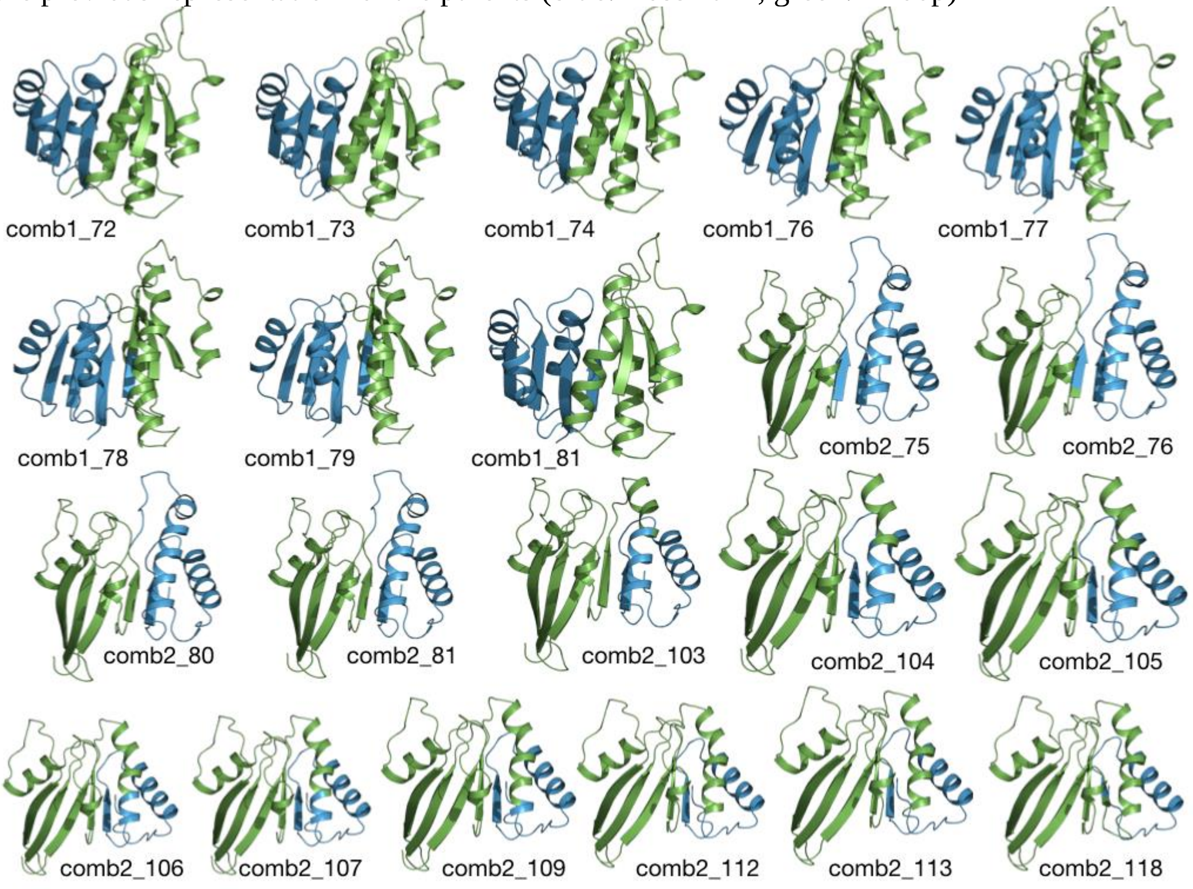

**Figure S9: The two largest hydrophobic clusters found in the parent domains and chimera comb1_72.** Largest and second largest clusters are depicted in black and white, respectively.

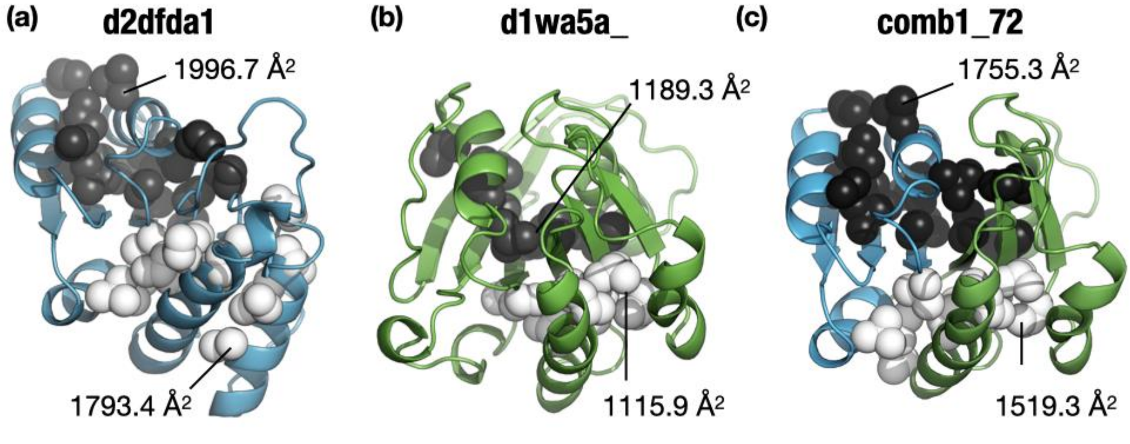

**Figure S10: Salt bridges for parent domains and chimera.** Acidic and basic residues are shown in red and blue, respectively.

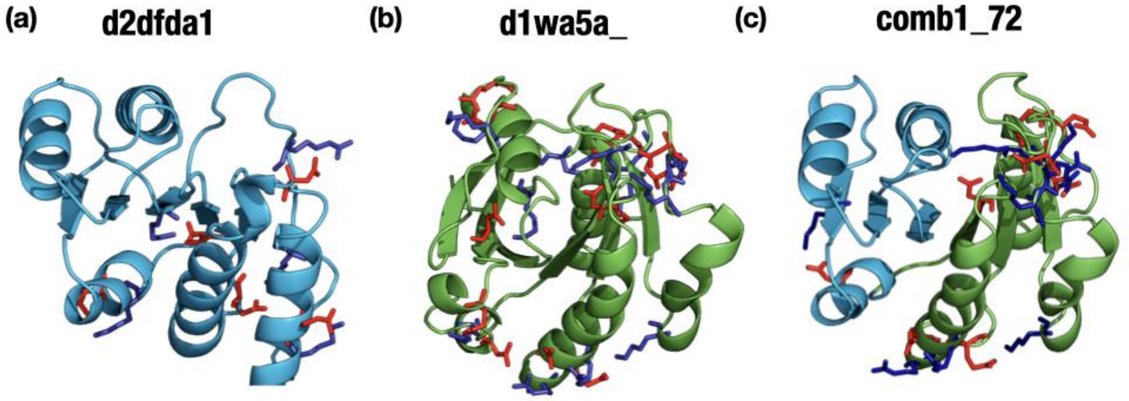

**Figure S11: Contact maps for parent domains and chimera.** Residues close in distance are shown in green, whereas those far apart are shown in different shades of blue. Other color representations are possible.

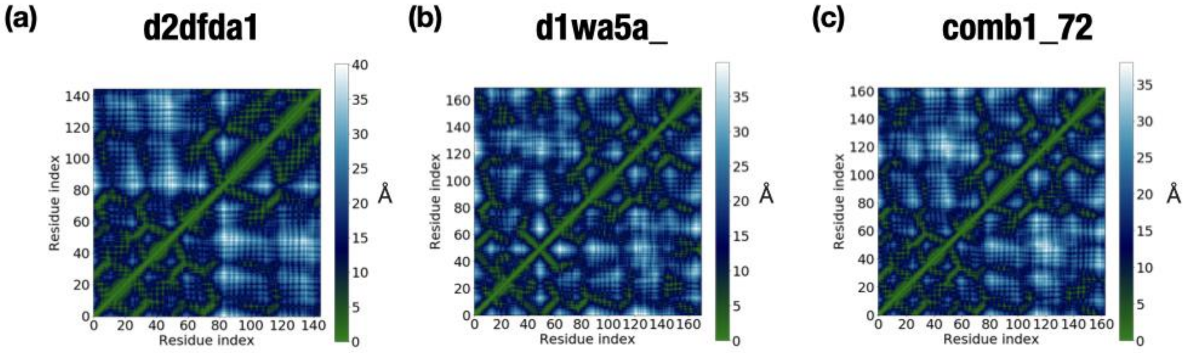
